# Mapping the Energy Landscape of PROTAC-Mediated Protein-protein Interactions

**DOI:** 10.1101/2021.08.31.458424

**Authors:** José A. Villegas, Tasneem M. Vaid, Michael E. Johnson, Terry W. Moore

## Abstract

One of the principal difficulties in computational modeling of macromolecules is the vast conformational space that arises out of large numbers of atomic degrees of freedom. This problem is a familiar issue in the area of protein-protein docking, where models of protein complexes are generated from the monomeric subunits. Although restriction of molecular flexibility is a commonly used approximation that decreases the dimensionality of the problem, the seemingly endless number of possible ways two binding partners can interact generally necessitates the use of further approximations to explore the search space. Recently, growing interest in using computational tools to build predictive models of PROTAC-mediated complexes has led to the application of state-of-the-art protein-protein docking techniques to tackle this problem. Additionally, the atomic degrees of freedom introduced by flexibility of linkers used in the construction of PROTACs further expands the configurational search space, a problem that can be tackled with conformational sampling tools. However, repurposing existing tools to carry out protein-protein docking and linker conformer generation independently results in extensive sampling of structures incompatible with PROTAC-mediated complex formation. Here we show that it is possible to restrict the search to the space of protein-protein conformations that can be bridged by a PROTAC molecule with a given linker composition by using a cyclic coordinate descent algorithm to position PROTACs into complex-bound configurations. We use this methodology to construct a picture of the energy landscape of PROTAC-mediated interactions in a model test case, and show that the global minimum lies in the space of native-like conformations.

## Introduction

The interaction between the proteome and the metabolome is possible as a result of the ability of folded proteins to harbor accessible compartments that can be occupied by guest molecules. Biochemical pathways proceed through these contact points by a variety of mechanisms, such as catalysis, allostery, and signal transduction across membranes. The treatment of disease at the molecular level has traditionally relied on the use of naturally derived or synthetic small molecules to suppress targeted pathways by blocking access to these contact points through competitive binding for the binding pocket. This approach requires high occupancy of the drug employed at the intended binding sites to effectively outcompete the targeted interaction. An alternative strategy that has recently gained attention is targeted protein degradation (TPD), in which the protein of interest is eliminated altogether by inducing its premature processing through the protein degradation machinery.^1^

E3 ligases are principal actors in the targeted degradation of proteins. The formation of a supra-molecular complex between the E3 ligase complex and a cytosolic protein results in catalyzed covalent bond formation between surface-exposed lysine residues and endogenously expressed ubiquitin proteins. The number of ubiquitin units appended, as well as the location of the attachment point, directs the protein toward specific protein degradation pathways.

Proteolysis targeting chimeras (PROTACS) are a novel class of drug candidates composed of two or more protein binding moieties (with at least one being an E3 ligase binder) joined together by a linker segment. Concurrent binding of the molecule to each of the targets is intended to bring the protein of interest in close proximity to the E3 ligase complex, resulting in its ubiquitination and subsequent degradation. It is clear that the linker is not a neutral actor in the formation of the supramolecular complex, as the linker lies at a central position between the interacting proteins. The linker must be capable of folding so that it does not interfere with the interaction between proteins, and must find a way to nestle at the protein-protein interface.^2^ Such folding should be energetically neutral at worst, but may also contribute to enhancing the protein-protein interaction by the formation of hydrophobic interactions at the protein complex surface.^3^

The difficulty in anticipating the three-dimensional arrangement of the resulting protein-PROTAC-protein complex makes the selection of the linker composition and length a non-trivial task. Typically this selection is carried out in an exploratory manner, where a library of candidate bifunctional molecules are synthesized with varying lengths and polymeric subunits.^4,5^ Accurate prediction of the structural determinants of PROTAC-mediated protein-protein interactions could serve to narrow the set of viable candidate molecules, saving valuable resources and shortening the drug-discovery timeline.

Modeling of PROTAC-bridged complexes encompasses two distinct problems, each of which covers vast areas of conformational space: protein-protein docking and linker conformational sampling.^6^ At the intersection between these two spaces exists a reduced subset of conformations where the protein-protein complex can be bridged by the bifunctional PROTAC molecule (Fig. 1). To date, strategies for the computational modeling of PROTAC-mediated complexes have attempted to explore each of these spaces independently and to identify from these searches a subset of structures existing at this intersection.

**Figure 1.**
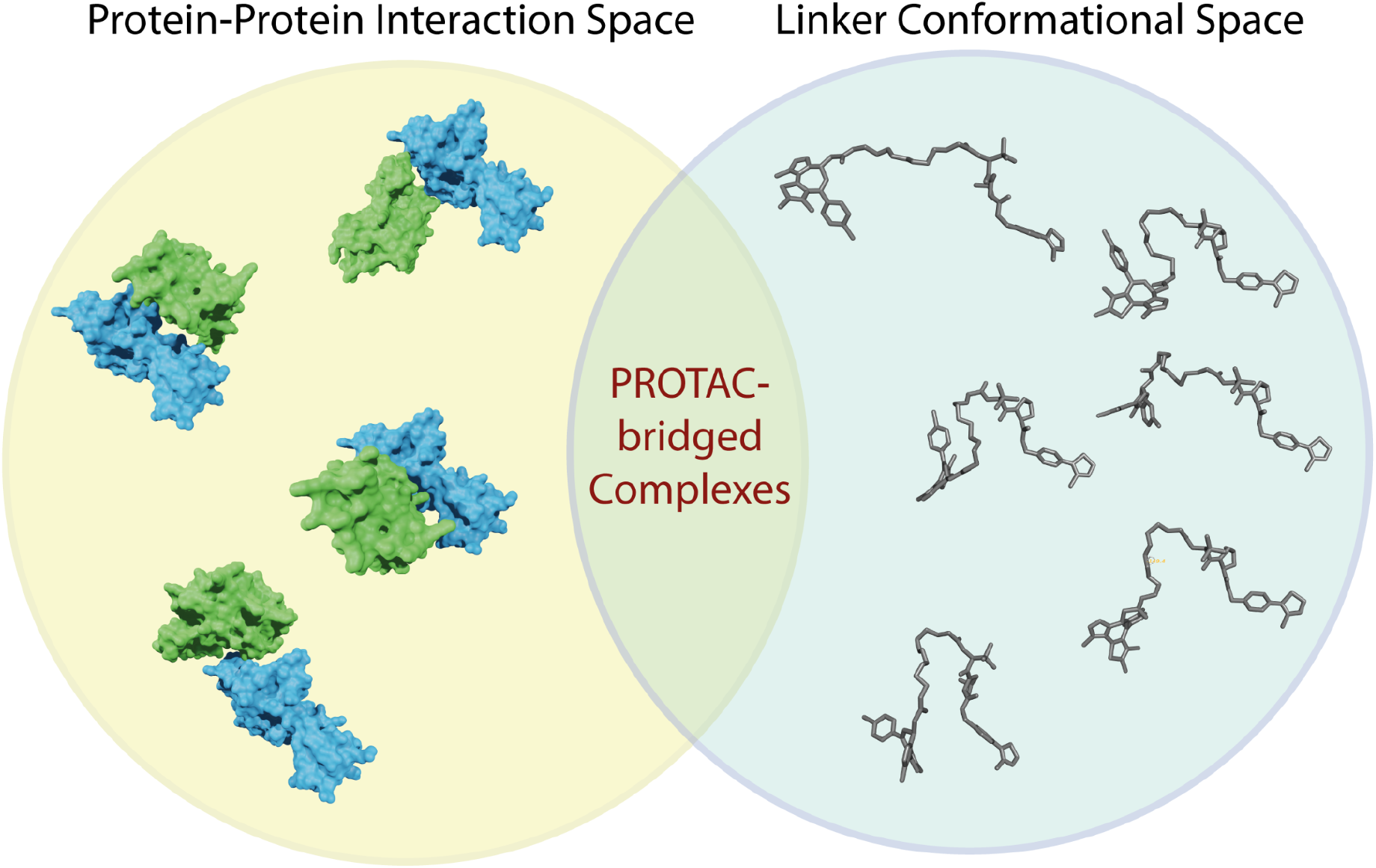
Schematic of the conformational search space in the modeling of PROTAC-bridged complexes. A large portion of the protein-protein interaction space is not compatible with PROTAC bridging (yellow sphere). Much of linker conformational space is not compatible with protein-PROTAC-protein complex formation. At the intersection of these two spaces lies the region of accessible states of PROTAC-bridged complexes.

In protein-protein docking calculations, the structure of the complex is predicted from the unbound subunits. In a typical setup, one molecule is kept fixed while the other is allowed to explore rotational and translational degrees of freedom. Within this six-dimensional search space there is a large number of conformations where the subunits are situated within van der Waals’ contact and can bury part of their surface areas. Efficient sampling of this space has necessitated the use of simplified representations, either in the form of discretization of a set of degrees of freedom, or by coarse-grained models of protein structure.^7^ Grid discretization enables the use of harmonic analysis methods to accelerate the search. In this approach, transformation of the representation into Fourier space allows for the direct calculation of the best displacement along chosen degrees of freedom according to the scoring function employed. In the coarse-grained model approach, protein structure is simplified to a reduced representation consisting of units which serve to model the collective behaviour of groups of atoms.^8^ As molecular interactions are calculated in an atom pairwise manner, a reduction in the number of “atoms” in the model results in an exponential decrease in computational cost. Simplified representations come at the cost of loss of structural information, and serve primarily to identify wide regions of protein-protein interaction space around the vicinity of the free energy minimum of interaction. This procedure, commonly referred to as a “global search”, is usually followed by a refinement step (“local search”) using a full atom representation.^7^

The second problem encountered in the modeling of PROTAC-bridged complexes is the conformational search along the torsional degrees of freedom of the linker. The high rotational freedom in commonly used linker chains results in large sets of possible conformers. Notably, Drummond and Williams used the PDB structure 5T35 to explore a number of different techniques for generation of linker conformations compatible with PROTAC-mediated protein-protein complex formation.^9,10^ In one methodology, the complex is placed in a configuration where the linker is fully extended, and displacements along linker rotational degrees of freedom are sampled with a Monte Carlo procedure. However, this method was found to perform poorly and was not able to recover any crystal-like poses (defined as an Cα RMSD ≤ 10 Å between the X-ray crystal structure coordinates and the predicted pose). The more successful methods relied on performing conformational sampling of the linker using MOE modelling software. The best performing method involved carrying out protein-protein docking and linker conformational sampling independently, and then combining the two sets of results to generate candidate ternary complexes. Bai *et. al*. followed a similar line of attack, where constrained protein-protein docking in Rosetta and linker conformational sampling using OMEGA were performed independently, and models were constructed by combining results from the two sets of data.^11^

Zaidman *et. al*. took a different approach, where data obtained from random sampling of PROTAC conformations was used to inform the protein-protein docking calculations.^12^ These data were fed in as constraints to an initial docking procedure, followed by refinement using unconstrained docking in RosettaDock. Generation of bridged complexes is attempted for each model using random conformational sampling of the linker.

Here, we develop a methodology to restrict the protein-protein docking procedure to the space of conformations where both partners are able to bind to the PROTAC molecule. The resulting space is greatly reduced relative to the ligand-free docking, since the linker can only bridge across proteins in a limited number of bound conformations. This constraint should allow for extensive sampling of the energy landscape of the accessible structures given a certain linker structure length. Exploration of this energy landscape would allow for the identification of regions of low-energy conformations.

To achieve this, we begin with a protein-PROTAC-protein complex with a fully extended linker. Each of the dihedral angles for rotatable bonds on the linker is set to be 180 degrees, and the proteins are placed appropriately to bind to their respective moieties (Fig. 2a). It may be that the proteins are placed in such a way that there is extensive overlap between the two molecules. However, these overlaps need not be a problem, since this high-energy conformation can be relaxed during the conformational sampling.^13^ When applying small torsional and translational displacements to the mobile chain, we can expect that the optimal linker conformation will not differ substantially from the previous step. Therefore, the conformational search along the linker rotational degrees of freedom can be restricted to deviate only within a certain range, reducing the space of linker conformations that must be explored.

**Figure 2.**
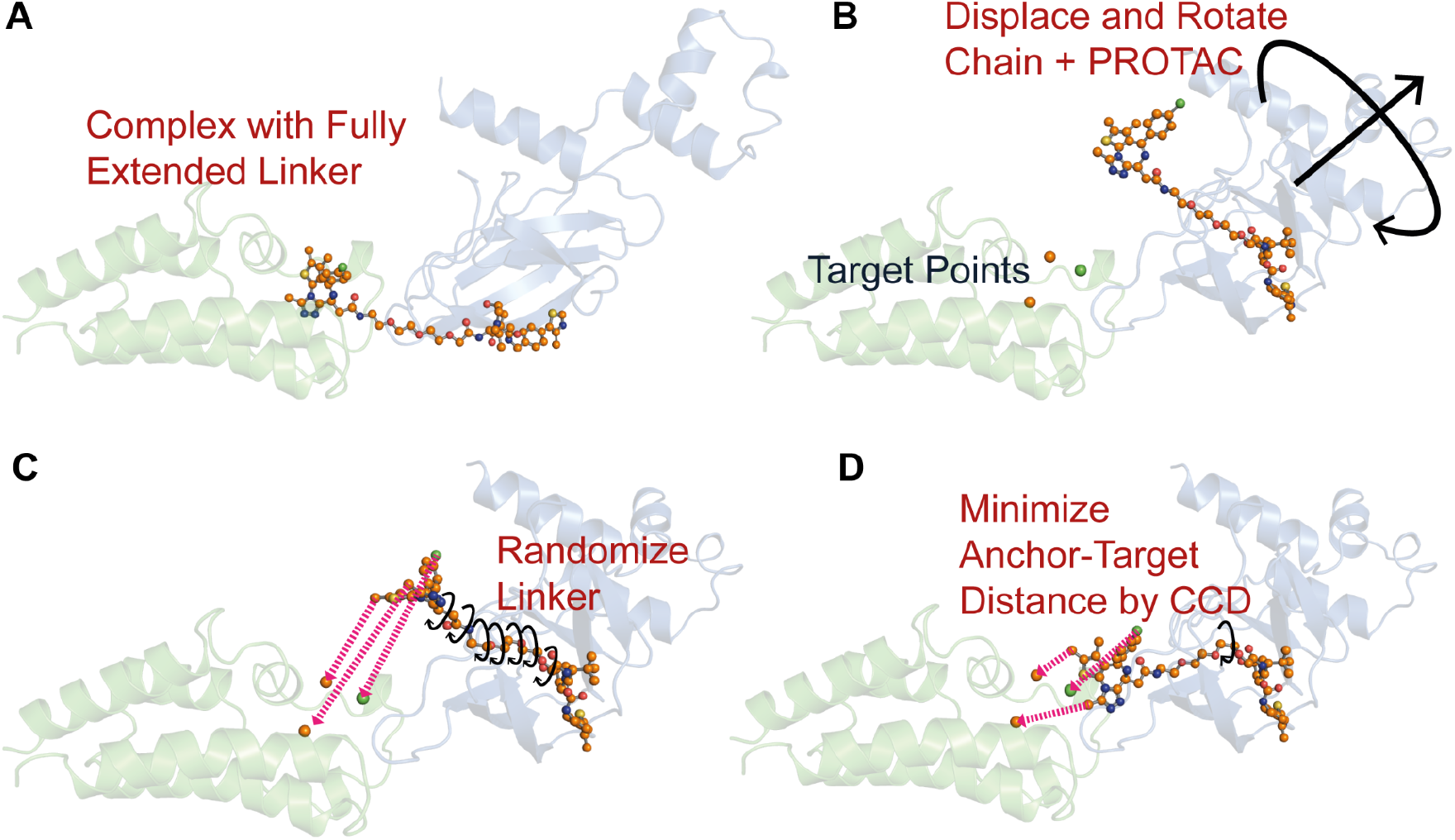
A) Initial configuration of protein-protein docking procedure. Linker is in fully extended conformation. B) Perturbation of mobile chain and PROTAC by random displacement along all six degrees of freedom. Target points are atomic coordinates of selected atoms on protein-binding moiety as found in the X-ray crystal structure file. C) Linker conformational variability is created by randomization of linker torsonal angles within selected bounds. D) PROTAC is repositioned in the fully bound configuration by rotation along linker rotatable bonds by using a cyclic coordinate descent algorithm.

In our methodology, the mobile chain is displaced along with the PROTAC molecule, and a Cyclic Coordinate Descent (CCD) procedure is employed to reposition the ligand back into the pocket of the stationary chain (Fig. 2b,d). In the process, an ensemble of linker conformations is generated by incorporating a randomization step (Fig. 2c). If a protein displacement is such that the linker is unable to bridge the two ligands, that displacement step is rejected. In this manner, the docking procedure is restricted to the space of bridgeable conformations.

## Methods

### Input Preparation

X-ray crystal structure coordinates were retrieved from the Protein Data Bank,^14^ and the protein-PROTAC-protein complex was prepared using the Protein Preparation Tool in Maestro (Schrodinger Release 2020-3). Hydrogens were added, the hydrogen bonding network was optimized, and protonation states were determined in the presence of the ligand. Structures were energy-minimized with heavy atoms constrained to within 0.3 Å. CHARMM22 topology and parameter files were constructed for the PROTAC ligands using the CGenFF web server.^15,16^

### PROTAC constrained protein-protein docking

Docking calculations were carried out by maintaining the E3 ligase protein stationary and applying rigid body transformations to the target-PROTAC complex. Torsional angles along the PROTAC linker region were adjusted by rotation of atoms on the E3 ligase-binding end of the molecule. To generate a starting conformation, all torsional angles along the linker were set to 180°, and target protein and PROTAC were displaced to align the E3 ligase binding moiety with the crystallographic position.

### Monte Carlo Sampling

Global Docking was carried out by using a simulated annealing procedure, where random rotations and translations of the target-PROTAC complex were performed. After each step in the search, the E3 binding moiety is displaced from its crystallographic position in the binding pocket. The conformation of the linker is therefore adjusted to reposition the E3 binding moiety at the correct location. This step is accomplished by using a cyclic coordinate descent (CCD) algorithm, as originally applied to protein loop closure calculations by Canutescu *et. al*.^17^

A range of linker conformations is considered by creating 100 copies of the PROTAC, and adjusting each of the linker torsional angles by a random amount before carrying out the CCD procedure. Complete randomization of the linker (assigning each torsional angle a value between 0° and 360°), yields conformations that are too distant from the starting conformation, resulting in a low success of linker closure. Therefore, we restrict randomization of each torsional angle to take on values of −135° to +135°. Although this reduction only restricts each torsional angle to 75% of its accessible value, it results in restricting the total linker conformational space to (0.75)^n^ of all possible values. For a linker with 14 rotatable bonds, only 1.78% of the total conformational space is accessed. By using the torsion from the previous step as the starting point for linker randomization, the search can be restricted to the space of linker conformations most accessible at the current step. Further randomization is obtained by selecting torsional angles at random during the CCD procedure, rather than sequentially as originally described.^17^ The RMSD criteria between the anchor and target atoms is set to 0.5 Å, and conformers that fail to close after 10,000 CCD steps are discarded.

Each conformer generated is assigned an initial probability, with all conformations being equally probable. Protein chains are treated using a rigid body approach, so that each side chain conformation is kept fixed and assigned a probability of 1.0. Scoring at each Monte Carlo step is done by using a probabilistic approach to calculate a Boltzmann weighted average potential energy over all conformations, with the conjugate parameter *β* set to 0.5 kcal/mol.^18–21^ This procedure allows for linker flexibility to be considered in the scoring, with multiple conformations contributing to the average energy. Potential energy terms were calculated using CHARMM22 vdW, electrostatic, and dihedral parameters. Protein intramolecular energy terms are ignored in the energy term, the cut-off for non-bonded interactions was set to 8 Å, and a cap of 1000 kcal/mol was placed on all pairwise energy terms. Points along the trajectory where no linker conformations are successfully generated are automatically rejected. A simulated annealing program was applied, with an exponential decay function.^22^ Runs consisted of 4000 sampling steps, starting from an effective temperature of *T_0_* =3000 and a decay constant of *τ_c_*=1000. At each MC trial, random displacements were applied to all six degrees of freedom, with maximum rotation step sizes set to 10° and maximum translation step sizes set to 0.1 Å.

## Results and Discussion

As has been noted by Bai *et. al*., modeling of PROTAC-mediated complexes should yield a complete picture of the accessible states in solution and not only aim to recover the X-ray crystal structure conformations. While molecules subjected to crystallization conditions anneal to a well-defined single state, the object of the PROTAC approach is to induce the association of protein partners which have not naturally evolved to form complexes. Therefore, it is unlikely these induced complexes exist in a single well-defined conformation in solution, as the resulting interfaces have not been optimized by nature. It is reasonable to assume that the free energy landscape should be composed of multiple energy minima, and lack a single deep energy well. Given the lack of a large enthalpic driver of protein-protein association, the presence of a number of accessible states would in theory help to lower the change in free energy between the bound and unbound forms, since there is an entropic benefit from having a set of accessible states rather than a single metastable state. We hypothesize that the ideal length of a PROTAC linker should not be one where the complex is not locked into one particular state, but rather the linker length should produce an energy landscape where the complex can form favorable interactions in a variety of conformations. However, the linker should not be so long as to create enthalpic and entropic penalties for the association of the PROTAC with the two binding partners.

To investigate the utility of our method in mapping the energy landscape of PROTAC mediated protein-protein interactions, we followed the approach of Drummond and Williams and used the PDB structure 5T35 as a test model. We considered chain H (Von Hippel-Lindau disease tumor suppressor) as the stationary chain, and chain E (Bromodomain-containing protein 4) as the mobile protein. Our objective was not only to recover the crystal structure of the native protein, but to map out the space of possible ternary complexes.

We therefore carried out an exhaustive search using a simulated annealing approach, starting from a fully extended PROTAC conformation. To determine if this procedure was capable of identifying a native-like conformation during a typical run, we carried out ten MC runs and selected the lowest energy conformation from each one. We observed that this methodology is capable of finding native-like poses with reliable reproducibility, where the lowest energy docked conformation has a Cα RMSD ≤ 10 Å in seven out of the Monte Carlo runs (Table 1.) The other three runs identified alternate conformations which were closely related. These poses represent an alternate conformation where favorable interactions can be made between the protein partners and the ligand.

| Replica | Energy (kcal/mol) | α-Carbon RMSD (Å) | Replica | Energy (kcal/mol) | α-Carbon RMSD (Å) |
| --- | --- | --- | --- | --- | --- |
| 1 | -252.204 | 7.5 | 6 | -239.256 | 9.8 |
| 2 | -228.692 | 24.4 | 7 | -264.127 | 5.5 |
| 3 | -249.892 | 24.9 | 8 | -288.955 | 7.3 |
| 4 | -247.938 | 24.8 | 9 | -267.819 | 6.2 |
| 5 | -250.945 | 9.4 | 10 | -260.176 | 7.6 |

We then aggregated the points sampled during all ten runs and used these to construct a full picture of the energy landscape using RMSD from the native conformation as the collective variable (Fig. 3A). A picture emerges of a landscape with two distinct minima, with the global minimum located in the region below the 10 Å cut-off point. The lowest energy structure has the mobile chain positioned in a nearly identical orientation as the crystal conformation, but with the two chains situated further apart (Fig. 3C). This is likely a result of the repulsive term of the vdW interaction becoming dominant at these close distances, preventing the complex from forming a tightly packed interface. Incorporation of an all-atom energy minimization step with backbone flexibility could potentially aid in resolving this over-sampling of interatomic overlaps and allow for closer contact between the subunits in the complex. However, native-like contacts are recovered in this model, with electrostatic interactions between the same amino acid side chain pairs (Fig. 3B).

**Figure 3.**
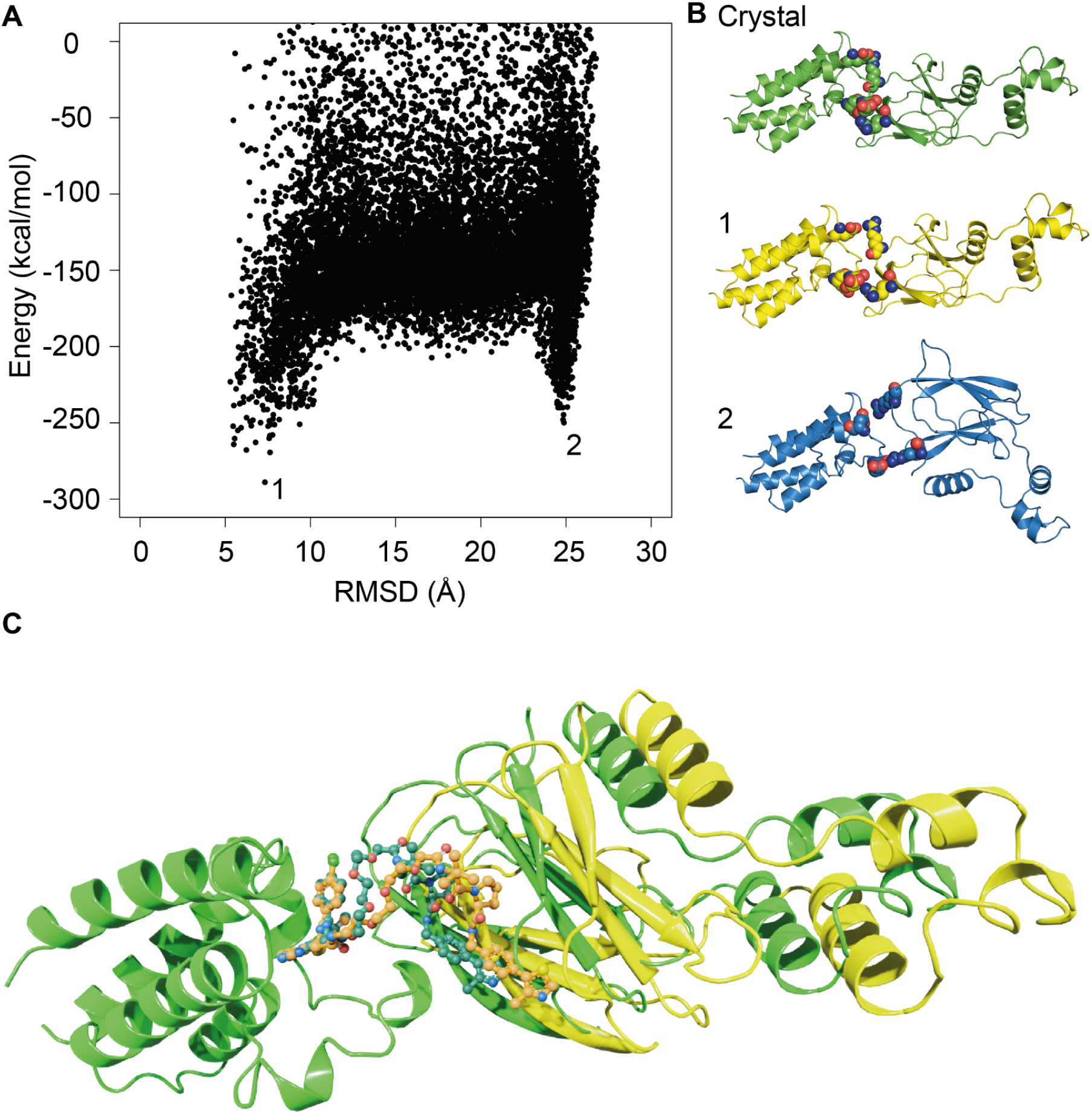
A) Energy landscape of protein-target interactions from 5T35 B) Ionizable residues at the protein-protein interface in the crystal structure and at energy landscape minima. C) Crystal structure of 5T35 (green). Lowest energy conformation sampled after ten Monte Carlo runs (yellow).

We also observe that the alternate conformations observed in three out of the ten Monte Carlo runs lie at a distinct local minimum. Although this minimum appears to be more shallow and narrow than the global minimum, the complexes in this region form stabilizing contacts along the resulting interface, and likely represent a conformational state which is significantly populated in solution.

Our methodology relies on choosing the starting point on the energy landscape where only one linker conformation is geometrically allowed. We are then able to efficiently build a set of PROTAC structures at each Monte Carlo step by assuming that the conformation of the linker should not deviate wildly from one step to the next. The ensemble of PROTAC structures at each step need not all have the same chemical composition. Candidates with different linker lengths and polymer subunits could potentially be generated. As linkers of different lengths will position the ternary complexes at different points on the landscape when fully extended, the design procedure would require traversing the landscape in such a way as to perform an initial pass through all these initial points. The capability to construct viable PROTAC-mediated complexes along the search trajectory opens up the possibility of introducing design capabilities into the procedure, with the potential to accelerate the process of designing bifunctional molecules for therapeutic purposes.

## Notes

### Competing Interest Statement

The authors have declared no competing interest.

## References

(1) Luh, L. M.; Scheib, U.; Juenemann, K.; Wortmann, L.; Brands, M.; Cromm, P. M. Prey for the Proteasome: Targeted Protein Degradation-A Medicinal Chemist’s Perspective. Angew.Chem. Int. Ed Engl. 2020, 59 (36), 15448–15466.

(2) Smith, B. E.; Wang, S. L.; Jaime-Figueroa, S.; Harbin, A.; Wang, J.; Hamman, B. D.; Crews, C. M. Differential PROTAC Substrate Specificity Dictated by Orientation of Recruited E3 Ligase. Nat. Commun. 2019, 10 (1), 131.

(3) Bondeson, D. P.; Smith, B. E.; Burslem, G. M.; Buhimschi, A. D.; Hines, J.; Jaime-Figueroa, S.; Wang, J.; Hamman, B. D.; Ishchenko, A.; Crews, C. M. Lessons in PROTAC Design from Selective Degradation with a Promiscuous Warhead. Cell Chem Biol 2018, 25 (1), 78–87.e5.

(4) Troup, R. I.; Fallan, C.; Baud, M. G. J. Current Strategies for the Design of PROTAC Linkers: A Critical Review. Exploration of Targeted Anti-tumor Therapy. 2020. https://doi.org/10.37349/etat.2020.00018.

(5) Zagidullin, A.; Milyukov, V.; Rizvanov, A.; Bulatov, E. Novel Approaches for the Rational Design of PROTAC Linkers. Exploration of Targeted Anti-tumor Therapy. 2020, pp 381–390. https://doi.org/10.37349/etat.2020.00023.

(6) Serapian, S. A.; Triveri, A.; Marchetti, F.; Castelli, M.; Colombo, G. Exploiting Folding and Degradation Machineries To Target Undruggable Proteins: What Can a Computational Approach Tell Us? ChemMedChem 2021, 16 (10), 1593–1599.

(7) Vakser, I. A. Protein-Protein Docking: From Interaction to Interactome. Biophys. J. 2014, 107 (8), 1785–1793.

(8) Kmiecik, S.; Gront, D.; Kolinski, M.; Wieteska, L.; Dawid, A. E.; Kolinski, A. Coarse-Grained Protein Models and Their Applications. Chem. Rev. 2016, 116 (14), 7898–7936.

(9) Drummond, M. L.; Williams, C. I. In Silico Modeling of PROTAC-Mediated Ternary Complexes: Validation and Application. J. Chem. Inf. Model. 2019, 59 (4), 1634–1644.

(10) Drummond, M. L.; Henry, A.; Li, H.; Williams, C. I. Improved Accuracy for Modeling PROTAC-Mediated Ternary Complex Formation and Targeted Protein Degradation via New In Silico Methodologies. J. Chem. Inf. Model. 2020, 60 (10), 5234–5254.

(11) Bai, N.; Miller, S. A.; Andrianov, G. V.; Yates, M.; Kirubakaran, P.; Karanicolas, J. Rationalizing PROTAC-Mediated Ternary Complex Formation Using Rosetta. J. Chem. Inf.Model. 2021, 61 (3), 1368–1382.

(12) Zaidman, D.; Prilusky, J.; London, N. PRosettaC: Rosetta Based Modeling of PROTAC Mediated Ternary Complexes. J. Chem. Inf. Model. 2020, 60 (10), 4894–4903.

(13) Allen, M. P.; Allen, M.; Tildesley, D. J.; Allen, T.; Tildesley, D. Computer Simulation of Liquids; Oxford University Press, 1989.

(14) Berman, H. M. The Protein Data Bank. Nucleic Acids Research. 2000, pp 235–242. https://doi.org/10.1093/nar/28.1.235.

(15) Vanommeslaeghe, K.; MacKerell, A. D. Automation of the CHARMM General Force Field (CGenFF) I: Bond Perception and Atom Typing. Journal of Chemical Information and Modeling. 2012, pp 3144–3154. https://doi.org/10.1021/ci300363c.

(16) Vanommeslaeghe, K.; Prabhu Raman, E.; MacKerell, A. D. Automation of the CHARMM General Force Field (CGenFF) II: Assignment of Bonded Parameters and Partial Atomic Charges. Journal of Chemical Information and Modeling. 2012, pp 3155–3168. https://doi.org/10.1021/ci3003649.

(17) Canutescu, A. A.; Dunbrack, R. L., Jr. Cyclic Coordinate Descent: A Robotics Algorithm for Protein Loop Closure. Protein Sci. 2003, 12 (5), 963–972.

(18) Kono, H.; Saven, J. G. Statistical Theory for Protein Combinatorial Libraries. Packing Interactions, Backbone Flexibility, and the Sequence Variability of a Main-Chain Structure. J. Mol. Biol. 2001, 306 (3), 607–628.

(19) Calhoun, J. R.; Kono, H.; Lahr, S.; Wang, W.; DeGrado, W. F.; Saven, J. G. Computational Design and Characterization of a Monomeric Helical Dinuclear Metalloprotein. J. Mol.Biol. 2003, 334 (5), 1101–1115.

(20) Bender, G. M.; Lehmann, A.; Zou, H.; Cheng, H.; Fry, H. C.; Engel, D.; Therien, M. J.; Blasie, J. K.; Roder, H.; Saven, J. G.; DeGrado, W. F. De Novo Design of a Single-Chain Diphenylporphyrin Metalloprotein. J. Am. Chem. Soc. 2007, 129 (35), 10732–10740.

(21) Fry, H. C.; Lehmann, A.; Sinks, L. E.; Asselberghs, I.; Tronin, A.; Krishnan, V.; Blasie, J. K.; Clays, K.; DeGrado, W. F.; Saven, J. G.; Therien, M. J. Computational de Novo Design and Characterization of a Protein That Selectively Binds a Highly Hyperpolarizable Abiological Chromophore. J. Am. Chem. Soc. 2013, 135 (37), 13914–13926.

(22) Zou, J.; Saven, J. G. Using Self-Consistent Fields to Bias Monte Carlo Methods with Applications to Designing and Sampling Protein Sequences. J. Chem. Phys. 2003, 118 (8), 3843–3854.

